# Integrase-associated niche differentiation of endogenous large DNA viruses in crustaceans

**DOI:** 10.1101/2023.01.31.526539

**Authors:** Satoshi Kawato, Reiko Nozaki, Hidehiro Kondo, Ikuo Hirono

## Abstract

Crustacean genomes harbor sequences originating from nimaviruses, a family of large double-stranded DNA viruses infecting crustaceans. In this study, we recovered metagenome-assembled genomes of 25 endogenous nimaviruses from crustacean genome data. Phylogenetic analysis revealed four major lineages within *Nimaviridae*, and for three of these lineages, we propose novel genera of endogenous nimaviruses: “Majanivirus” and “Pemonivirus” identified from penaeid shrimp genomes, and “Clopovirus” identified from terrestrial isopods. Majanivirus genomes contain multiple eukaryotic-like genes such as baculoviral inhibitor of apoptosis repeat-containing genes, innexins, and heat shock protein 70-like genes, some of which contain introns. An alignment of long reads revealed that that each endogenous nimavirus species specifically inserts into host microsatellites or within 28S rDNA. This insertion preference was associated with the type of virus-encoded DNA recombination enzymes, the integrases. Majaniviruses, pemoniviruses, some whispoviruses, and possibly clopoviruses specifically insert into the arthropod telomere repeat motif (TAACC/GGTTA)n and all possessed a specific tyrosine recombinase family. Pasiphaea japonica whipovirus and Portunus trituberculatus whispovirus, the closest relatives of white spot syndrome virus, integrate into the host 28S rDNA and are equipped with members of another family of tyrosine recombinases that are distantly related to telomere-specific tyrosine recombinases. Endogenous nimavirus genomes identified from sesarmid crabs, which lack tyrosine recombinases and are flanked by a 46-bp inverted terminal repeat, integrate into (AT/TA)n microsatellites through the acquisition of a Ginger2-like cut-and-paste DDE transposase. These results suggest that endogenous nimaviruses are giant transposable elements that occupy different sequence niches through the acquisition of different integrase families.

**Importance:** Crustacean genomes harbor sequences originating from a family of large DNA viruses called nimaviruses, but it is unclear why they are present. We show that endogenous nimaviruses selectively insert into repetitive sequences within the host genome, and this insertion specificity was correlated with different types of integrases, which are DNA recombination enzymes encoded by the nimaviruses themselves. This suggests that endogenous nimaviruses have colonized various genomic niches through the acquisition of integrases with different insertion specificities. Our results point to a novel survival strategy of endogenous large DNA viruses colonizing the host genomes. These findings may clarify the evolution and spread of nimaviruses in crustaceans and lead to measures to control and prevent the spread of pathogenic nimaviruses in aquaculture settings.

## Introduction

*Nimaviridae* is a family of double-stranded DNA viruses infecting crustaceans (1). The only officially recognized member, white spot syndrome virus (WSSV), is the most devastating viral pathogen affecting global shrimp aquaculture (1–4). Although several other crustacean viruses have been reported to exhibit morphological characteristics similar to those of nimaviruses (5, 6), only one virus, Chionoecetes opilio bacilliform virus (CoBV), has been verified at the sequence level (NCBI Accession no. BDLS01000001-BDLS01000002, LC741431).

Despite the limited number of known exogenous nimaviruses, genomic analyses of decapod crustaceans have revealed the presence of sequences originating from nimaviruses (7–11). These endogenous viral elements (12, 13) are present as multi-copy elements sometimes reaching hundreds of copies per haploid genome (7, 10). However, the biological significance of these endogenous nimaviruses is unknown, and they do not exhibit any virulence.

In this study, we reconstructed 24 complete genomes and one partial genome of endogenous nimaviruses recovered from crustacean genome data. Our results indicate that these viruses preferentially integrate into specific motifs in the host genome, and that this insertion specificity is tightly linked with the presence of different integrase-like enzymes encoded by the viral genomes. These observations suggest that endogenous nimaviruses are selfish genetic elements that have colonized the crustacean genomes.

## Results

### Metagenome-assembled genomes (MAGs) of endogenous nimaviruses

Genome sequencing of 17 crustacean genomes yielded 19 to 25 gigabases (Gb) of Illumina reads per genome and 769 Mb to 16.8 Gb of ONT reads per genome (Supplementary Table 1). We also analyzed publicly available sequence data of the swimming crab *Portunus trituberculatus* (14) and the terrestrial isopod *Trachelipus rathkii* (15).

Our analysis yielded a total of 25 endogenous nimaviral genomes, 24 of which were regarded as complete (Table 2). Of these, 23 genomes were deposited in the DDBJ/NCBI/ENA databases as metagenome assembled genomes (MAGs) of uncultivated virus genomes. The genomes of Portunus trituberculatus whispovirus and Trachelipus rathkii clopovirus are available as supplementary files of the manuscript (Supplementary File 1-4). These MAGs are consensus sequences of closely related clones infecting a single organism. Most of the coding sequences on the assembled genomes are intact, but the actual individual copies within the host genome may be disrupted by mutations. Despite these limitations, the MAGs of endogenous nimaviruses represent distinct lineages of nimaviruses and provide valuable information for analyzing the evolution of *Nimaviridae*.

**Table 1.** Crustacean samples sequenced in this study.

| **Genus** | **Species** | **Isolate** | **Geographic loaction** | **Year** | **BioSample Accession** |
| --- | --- | --- | --- | --- | --- |
| *Marsupenaeus* | *japonicus* | Aichi2020 | Japan: Aichi | 2020 | SAMD00511450 |
| *Melicertus* | *latisulcatus* | Mellat | Japan: Okinawa, East China Sea | 2016 | SAMD00111282 |
| *Penaeus* | *monodon* | Penmon | Japan: Aichi, Mikawa Bay | 2016 | SAMD00111283 |
| *Penaeus* | *semisulcatus* | Kagawa2020 | Japan: Kagawa | 2020 | SAMD00511443 |
| *Litopenaeus* | *vannamei* | Litvan | Japan | 2015 | SAMD00111285 |
| *Metapenaeus* | *ensis* | Metens | Japan: Aichi, Mikawa Bay | 2015 | SAMD00111287 |
| *Metapenaeus* | *joyneri* | Tokushima2020 | Japan: Tokushima | 2020 | SAMD00511444 |
| *Metapenaeopsis* | *lamellata* | Hokkoku2021 | Japan | 2021 | SAMD00511449 |
| *Trachysalambria* | *curvirostris* | Ube2021 | Japan: Yamaguchi, Ube | 2021 | SAMD00511448 |
| *Sicyonia* | sp. | Fukuoka2019 | Japan: Fukuoka | 2019 | SAMD00513257 |
| *Pasiphaea* | *japonica* | Toyama2020 | Japan: Toyama, Toyama Bay | 2020 | SAMD00511445 |
| *Hemigrapsus* | *takanoi* | Hemtak | Japan: Tokyo, Tokyo Bay | 2016 | SAMD00111291 |
| *Orisarma (Sesarmops)* | *intermedium* | Sesint | Japan: Kochi | 2016 | SAMD00111292 |
| *Orisarma (Chiromantes)* | *dehaani* | Chideh | Japan: Kochi | 2016 | SAMD00111293 |
| *Armadillidium* | *vulgare* | TUMSAT20210906 | Japan: Tokyo | 2021 | SAMD00511446 |
| *Porcellio* | *scaber* | TUMSAT20211004 | Japan: Tokyo | 2021 | SAMD00511447 |

**Table 2.** Nimaviral genomes characterized in this study.

| Proposed genus | Species | Host | Abbreviation | Accession | Length | GC% | CDS | Completeness | Coverage |  |
| --- | --- | --- | --- | --- | --- | --- | --- | --- | --- | --- |
|  |  |  |  |  |  |  |  |  | Illumina | ONT |
| "Majanivirus" | Marsupenaeus japonicus endogenous nimavirus | *Marsupenaeus japonicus* | MjeNMV | LC738868.1 | 306,008 | 33 | 111 | Complete | 23466.0 | 615.1 |
|  | Melicertus latisulcatus majanivirus | *Melicertus latisulcatus* | MelaMJNV | LC738874.1 | 287,061 | 32 | 104 | Complete | 769.5 | 173.7 |
|  | Metapenaeopsis lamellata majanivirus | *Metapenaeopsis lamellata* | MellatMJNV | AP027153.1 | 267951 | 27 | 106 | Complete | 118.9 | 6.6 |
|  | Penaeus monodon majanivirus A | *Penaeus monodon* | PemoMJNVA | LC738870.1 | 294144 | 40 | 115 | Complete | 649.4 | 69.4 |
|  | Penaeus monodon majanivirus B | *Penaeus monodon* | PemoMJNVB | LC738871.1 | 360,489 | 35 | 114 | Complete | 325.9 | 22.4 |
|  | Litopenaeus vannamei majanivirus Nimav-Lv_1 | *Litopenaeus vannamei* | LvMJNV | LC738872.1 | 280452 | 35 | 119 | Complete | 116.7 | 8.3 |
|  | Penaeus semisulcatus majanivirus | *Penaeus semisulcatus* | PeseMJNV | LC738873.1 | 291934 | 42 | 110 | Complete | 469.4 | 45.2 |
|  | Metapenaeus ensis majanivirus | *Metapenaeus ensis* | MeenMJNV | LC738876.1 | 292,272 | 29 | 101 | Complete | 207.1 | 17.0 |
|  | Metapenaeus joyneri majanivirus | *Metapenaeus joyneri* | MejoMJNV | LC738878.1 | 401182 | 26 | 124 | Complete | 169.8 | 17.9 |
|  | Trachysalambria curvirostris majanivirus | *Trachysalambria curvirostris* | TrcuMJNV | LC738879.1 | 283150 | 28 | 101 | Complete | 134.2 | 14.7 |
| "Pemonivirus" | Marsupenaeus japonicus pemonivirus | *Marsupenaeus japonicus* | MjPMNV | AP027202.1 | 323944 | 48 | 102 | Complete | 1187.1 | 4.7 |
|  | Melicertus latisulcatus pemonivirus | *Melicertus latisulcatus* | MelaPMNV | LC738875.1 | 359647 | 48 | 109 | Complete | 212.6 | 41.6 |
|  | Penaeus monodon endogenous nimavirus | *Penaeus monodon* | PmeNMV | LC738869.1 | 300002 | 45 | 94 | Complete | 892.6 | 95.2 |
|  | Penaeus semisulcatus pemonivirus | *Penaeus semisulcatus* | PesePMNV | AP027152.1 | 277334 | 43 | 102 | Complete | 157.0 | 6.5 |
| "Whispovirus" | Hemigrapsus takanoi nimavirus | *Hemigrapsus takanoi* | HtINMV | LC738882.1 | 251731 | 47 | 111 | Partial | 383.7 | 19.4 |
|  | Trachysalambria curvirostris nimavirus | *Trachysalambria curvirostris* | TcNMV | LC738880.1 | 331684 | 47 | 107 | Complete | 706.6 | 41.8 |
|  | Sesarmops intermedium nimavirus | *Sesarmops intermedium* | SiNMV | LC738884.1 | 267936 | 44 | 104 | Complete | 79.1 | 50.2 |
|  | Chiromantes dehaani nimavirus | *Chiromantes dehaani* | CdNMV | AP027155.1 | 285096 | 44 | 99 | Complete | 53.7 | 30.2 |
|  | Metapenaeus ensis nimavirus | *Metapenaeus ensis* | MeNMV | LC738877.1 | 341,283 | 44 | 117 | Complete | 517.7 | 55.1 |
|  | Sicyonia whispovirus | *Sicyonia* sp. | SicyWSV | LC738881.1 | 347,493 | 54 | 89 | Complete | 1351.1 | 237.7 |
|  | Portunus trituberculatus whispovirus | *Portunus trituberculatus* | PotrWSV | NA* | 323740 | 48 | 98 | Complete | 102.1 | 72.5 |
|  | Pasiphaea japonica whispovirus | *Pasiphaea japonica* | PajaWSV | LC738885.1 | 276272 | 35 | 86 | Complete | 159.4 | 32.4 |
| "Clopovirus" | Armadillidium vulgare clopovirus | *Armadillidium vulgare* | AvCLPV | LC738883.1 | 416,069 | 32 | 120 | Complete | 315.6 | 78.0 |
|  | Porcellio scaber clopovirus | *Porcellio scaber* | PsCLPV | AP027154.1 | 509411 | 31 | 179 | Complete | 1224.2 | 99.5 |
|  | Trachelipus rathkii clopovirus | *Trachelipus rathkii* | TrCLPV | NA* | 579413 | 35 | 196 | Complete | 125.4 | 112.8 |

\*Not applicable to third party annotation (TPA)

Maximum phylogenetic analysis of nimaviral core proteins (10, 16) revealed four major clusters within *Nimaviridae* (Figure 1). As discussed below, we believe that these clades represent distinct genus-level taxa.

**Figure 1.**
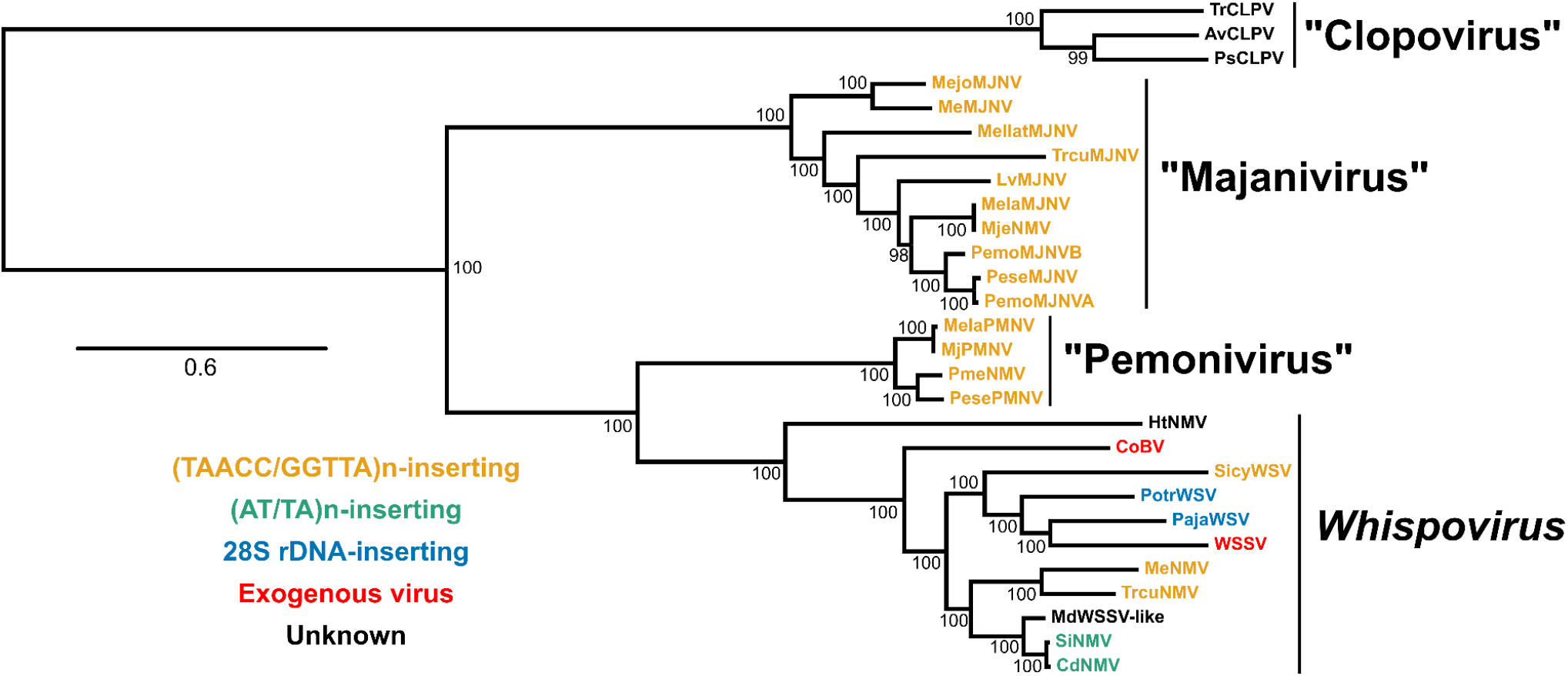
Phylogenomic tree of *Nimaviridae*. Amino acid sequences of nine nimaviral core genes (wsv026, wsv282, wsv289, wsv303, wsv343, wsv360, wsv433, wsv447, and wsv514; 13,007 amino acids) were used for the analysis. See Table 2 for the abbreviations.

### Majaniviruses colonizie telomere repeats of penaeid shrimp genomes

We previously reported on a group of penaeid shrimp-specific endogenous nimaviruses, exemplified by Marsupenaeus japonicus endogenous nimavirus (MjeNMV; LC738868.1) (Figure 2A) (10). We propose for these penaeid endogenous nimaviruses a genus-level cluster, “Majanivirus” (**<u>Ma</u>**rsupenaeus **<u>ja</u>**ponicus endogenous **<u>ni</u>**mavirus), consisting of 11 members. Majaniviral genomes range from 278 kb to 401 kb in size, with GC content ranging from 27% to 42%. Complete majanivirus genomes were recovered from all penaeid shrimp genome data except for *Sicyonia* sp. Kyushu2019.

**Figure 2.**
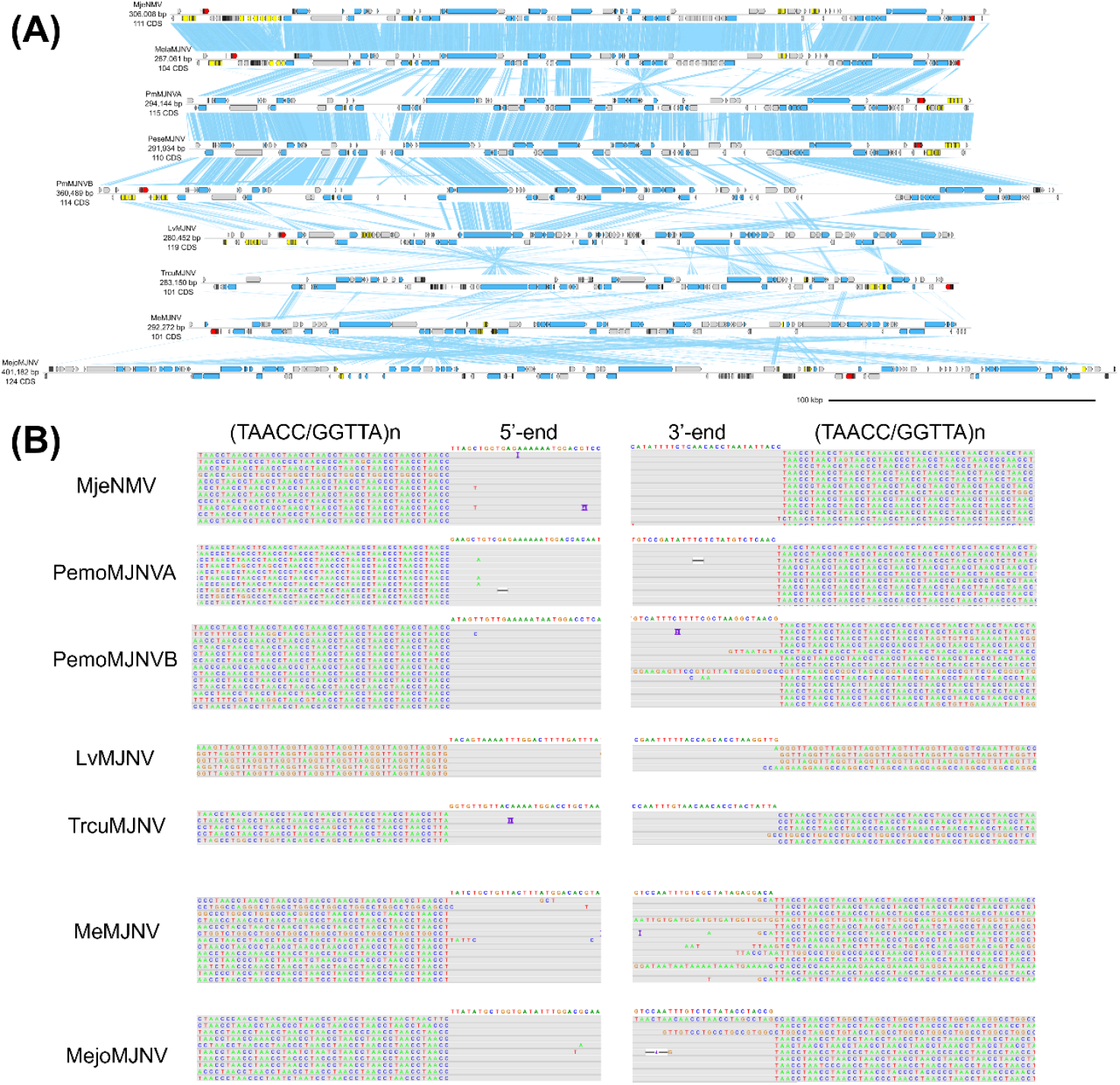
Genome diagram of Majaniviruses. (A) Genome diagrams of majaniviruses. Arrows indicate predicted genes and their transcriptional orientations; blue, WSSV homologs; yellow, baculoviral inhibitor of apoptosis repeat-containing proteins; gray: hypothetical and other eukaryotic-like proteins; red, tyrosine recombinases. Blue ribbons indicate pairwise TBLASTX hits (e-value :1-e3, bitscore: 50). (B) ONT read alignments flanking the 5’- and 3’-ends of maajnivirus genomes.

We previously showed that the MjeNMV genome is chromosomally integrated into the kuruma shrimp genome (10), but their integration sites remained unknown. Bao *et al*. (2020) were the first to show that Nimav-1_LVa (LC738872.1), a majanivirus, specifically insert into the (TAACC/GGTTA)n motifs in the genome of the Pacific white shrimp *Litopenaeus vannamei* (11). Our analysis of ONT read alignments indicate that MjeNMV and other complete majanivirus genomes are flanked by the same (GGTTA/TAACC)n pentanucleotide motifs (Figure 2B), strongly suggesting that telomere insertion is a common feature of majaniviruses. However, some ONT reads from one end of the (GGTTA/TAACC)n tract to the other end of the majaniviral genomes could be aligned, suggesting that some majaniviral copies are present as concatemers and/or episomes. This suggests that they are, or were until recently, actively replicating within the host genome.

A salient feature of majanivirus genomes is the expansion of eukaryotic-like genes (Figure 3). The earliest reportson WSSV-like sequences in the penaeid shrimp genomes noted an expansion of a large DNA segment containing WSSV homologs as well as various eukaryotic genes, including baculoviral inhibitor of apoptosis repeat (BIR)-containing proteins (7) and an HSP70 homolog (17). Bao et al. also observed the presence of eukaryotic-like genes on the Nimav-1_LVa genome (11). The availability of complete majaniviral genomes confirms that the presence of eukaryotic-like genes is a shared trait of majaniviruses. Heat shock protein 70-like proteins (MjHSP70-2) (17) and innexins form their own clades on the phylogenetic trees, indicating that they have been vertically inherited from a common ancestor of the majaniviruses (Figure 3). BIR-containing proteins clustered with other decapod proteins, but we surmise that they are nimaviral sequences annotated as host genes. These findings demonstrate that majaniviruses harbor multiple eukaryotic-like genes, which were likely acquired from their decapod hosts.

**Figure 3.**
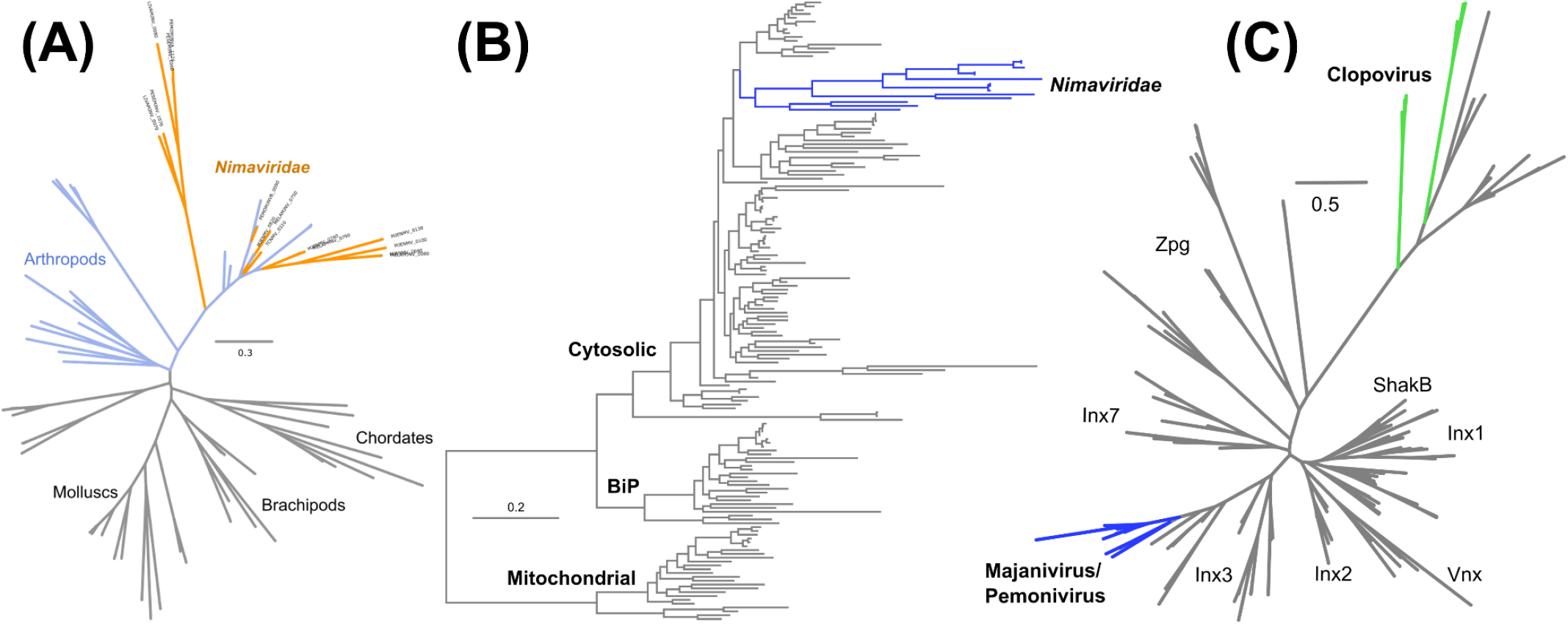
Phylogenetic analysis of eukaryotic-like genes in majaniviruses. (B) Maximum-likelihood phylogenetic tree of 80 baculoviral inhihbitor of apoptosis repeat-containing proteins (266 sites; model: LG+I+G4). (B) Maximum-likelihood phylogenetic tree of 162 HSP70-like proteins (579 sites; model: LG+R7). (C) Maximum-likelihood phylogenetic tree of 158 innexin proteins (251 sites; model: LG+R6).

Nimaviral core genes are a set genes that are ubiquitously conserved among *Nimaviridae* and are likely to play essential functions in the viral replication cycle (10, 11, 16). The original nimaviral core gene set consisted of 28 genes. Bao et al. proposed the inclusion of four additional genes (wsv112, wsv206, wsv226, wsw308) to this set, raising the total number to 32 (11). Our analysis supported the inclusion of wsv226, wsv308, and wsv310 as core genes, but phylogenetic analysis of wsv112 and wsv206 suggested that they were acquired independently (Supplementary Figure 1), although this does not necessarily mean that their functions are dispensable for viral replication. Consequently, our revised version of the nimaviral core gene set includes 31 genes (Table 3).

**Table 3.** Nimaviral core genes found in clopovirus genomes.

| Nimaviral core genes |  | ID | AvCLPV | PsCLPV | TrCLPV |
| --- | --- | --- | --- | --- | --- |
| Structural proteins | Envelope proteins | wsv293a |  |  |  |
|  |  | wsv327* | BDT63349.1 | BDV50114.1 | TRCLPV_1850 |
|  | Capsid proteins | wsv037 |  |  |  |
|  |  | wsv220 |  |  |  |
|  |  | wsv271 |  |  |  |
|  |  | wsv289 | BDT63373.1 | BDV50138.1 | TRCLPV_2110 |
|  |  | wsv308 |  |  |  |
|  |  | wsv332 |  |  |  |
|  |  | wsv360 | BDT63312.1 | BDV50077.1 | TRCLPV_0400 |
|  |  | wsv415 |  |  |  |
|  | Unknown | wsv131 |  |  |  |
|  |  | wsv161 |  |  |  |
| Nonstructural proteins | DNA polymerase | wsv514 | BDT63304.1 | BDV50068.1 | TRCLPV_0300 |
|  | Helicase | wsv447 | BDT63392.1 | BDV49995.1 | TRCLPV_1500 |
|  | AAA ATPase | wsv026 | BDT63390.1 | BDV49990.1 | TRCLPV_1520 |
|  | Protein kinase | wsv423 |  |  |  |
|  | Latency-related gene | wsv427 |  |  |  |
|  | TATA-binding protein | wsv303 | BDT63332.1 | BDV50131.1 | TRCLPV_2040 |
|  | hypothetical protein | wsv137 |  |  |  |
|  |  | wsv139 |  |  |  |
|  |  | wsv192 |  |  |  |
|  |  | wsv133 |  |  |  |
|  |  | wsv267 |  |  |  |
|  |  | wsv282 | BDT63296.1 | BDV50018.1 | TRCLPV_0140 |
|  |  | wsv285 |  |  |  |
|  |  | wsv310 |  |  |  |
|  |  | wsv313 |  |  |  |
|  |  | wsv343 | BDT63282.1 | BDV50024.1 | TRCLPV_1200 |
|  |  | wsv433 | BDT63379.1 | BDV50142.1 | TRCLPV_1580 |
|  |  | wsv440 |  |  |  |
| Missing in some<br>nimaviruses | Envelope proteins | wsv011* | BDT63370.1 | BDV50135.1 | TRCLPV_2140 |
|  |  | wsv021 | BDT63369.1 |  | TRCLPV_2140 |
|  |  | wsv035* | BDT63331.1 | BDV50132.1 | TRCLPV_2070 |
|  |  | wsv115* | BDT63386.1 | BDV49987.1 | TRCLPV_1560 |
|  |  | wsv209* | BDT63346.1 | BDV50116.1 | TRCLPV_1880 |
|  |  | wsv206 |  |  |  |
|  |  | wsv216 | BDT63368.1 | BDV50096.1 | TRCLPV_0600 |
|  |  | wsv325 | BDT63338.1 | BDV50126.1 | TRCLPV_1970 |
|  |  | wsv432 |  |  |  |
|  | Tegument | wsv306* | BDT63353.1 | BDV50110.1 | TRCLPV_1810 |
|  | Unknown | wsv134* | BDT63382.1 | BDV49996.1 | TRCLPV_1490 |
|  |  | wsv136 |  |  |  |

\*Naldaviral core genes

### Pemoniviruses: another telomere-dwelling endogenous nimavirus lineage colonizing penaeid shrimp genomes

We reconstructed the genomes of Penaeus monodon endogenous nimavirus (PmeNMV; LC738869.1) and three related viruses, forming a novel genus-level clade which we have named Pemonivirus (**<u>Pe</u>**naeus **<u>mo</u>**nodon **<u>ni</u>**mavirus; Figure 4A). Pemonivirus genomes range in size from 300 kb to 360 kb, with GC contents ranging from 43% to 48%. Pemoniviruses selectively insert into telomere motifs (Figure 4B). Unlike majaniviruses, pemonivirus genomes contain few eukaryotic-derived genes. Pemoniviruses were found only from members of Penaeus *sensu lato* occurring in the Indo-Western Pacific region, including *Marsupenaeus japonicus, Melicertus latisulcatus, Penaeus semisulcatus*, and *Penaeus monodon*.

**Figure 4.**
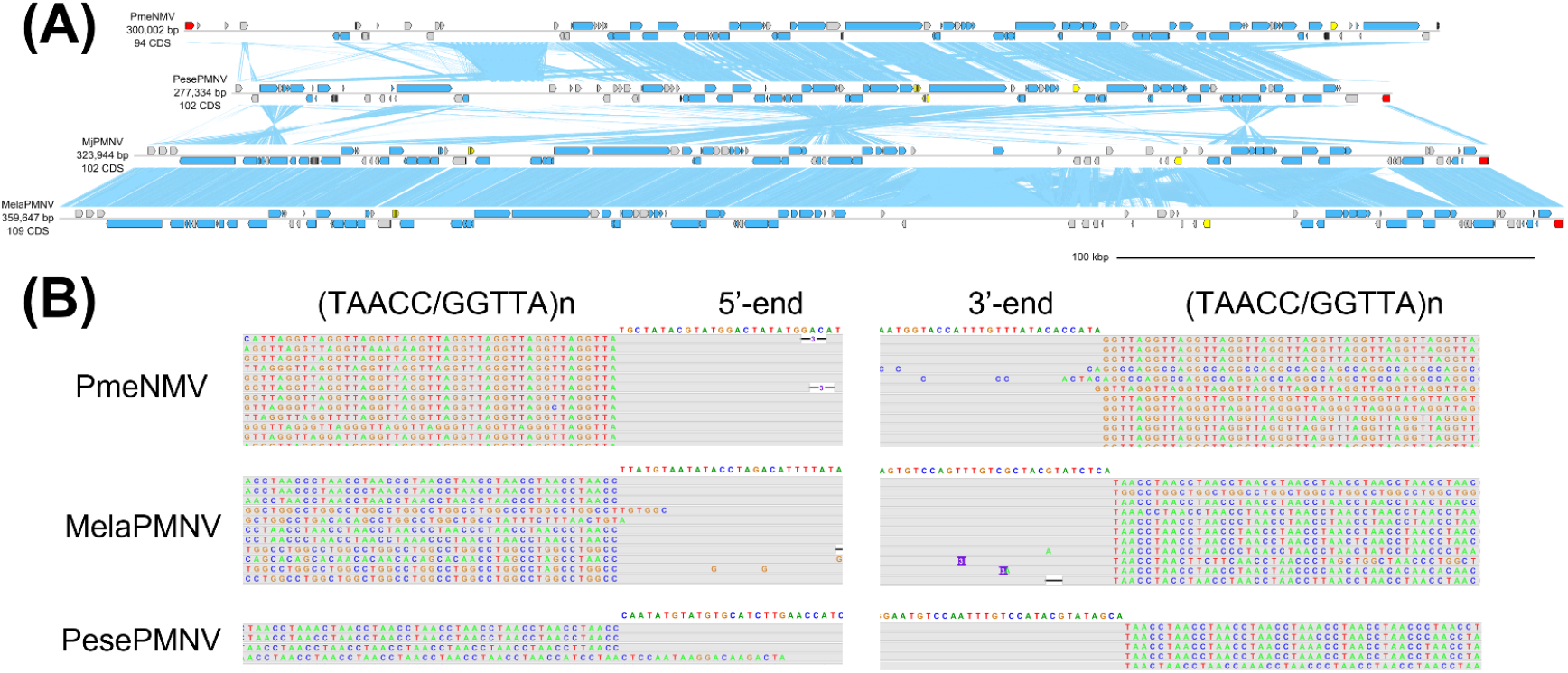
Genome diagrams of Pemoniviruses. (A) Genome diagrams of pemoniviruses. Arrows indicate predicted genes and their transcriptional orientations; blue, WSSV homologs; yellow, baculoviral inhibitor of apoptosis repeat-containing proteins; gray: hypothetical and other eukaryotic-like proteins; red, tyrosine recombinases. Blue ribbons indicate pairwise TBLASTX hits (e-value :1-e3, bitscore: 50) (B) ONT read alignments flanking the 5’- and 3’-ends of pemonivirus genomes.

### Clopovirus: divergent, terrestrial isopod-associated nimaviruses

We identified a new clade of nimaviruses in the genomes of terrestrial isopods, which we named Clopovirus (derived from **<u>clopo</u>**rte, the French word for “pill bug”) (Figure 5). Clopovirus genomes range in size from 410 kb to 580 kb, making them the largest nimavirus genomes discovered to date. Armadillidium vulgare clopovirus (AvCLPV; LC738883.1) was assembled into a 416,069 bp sequence, coding for 120 protein-coding genes. AvCLPV might be the same virus as the WSSV-like sequences reported by Thézé et al. (21). Porcellio scaber clopovirus (PsCLPV) was identified from the shotgun sequencing data of *Porcellio scaber*. The genome sequence of Trachelipus rathkii clopovirus (TrCLPV; Supplementary Files 1 and 2) was identified from the shotgun sequencing data of *Trachelipus rathkii*, a species native to Europe but introduced into North America (15). TrCLPV has a genome size of 579 kb, the largest of all nimaviruses discovered so far.

**Figure 5.**
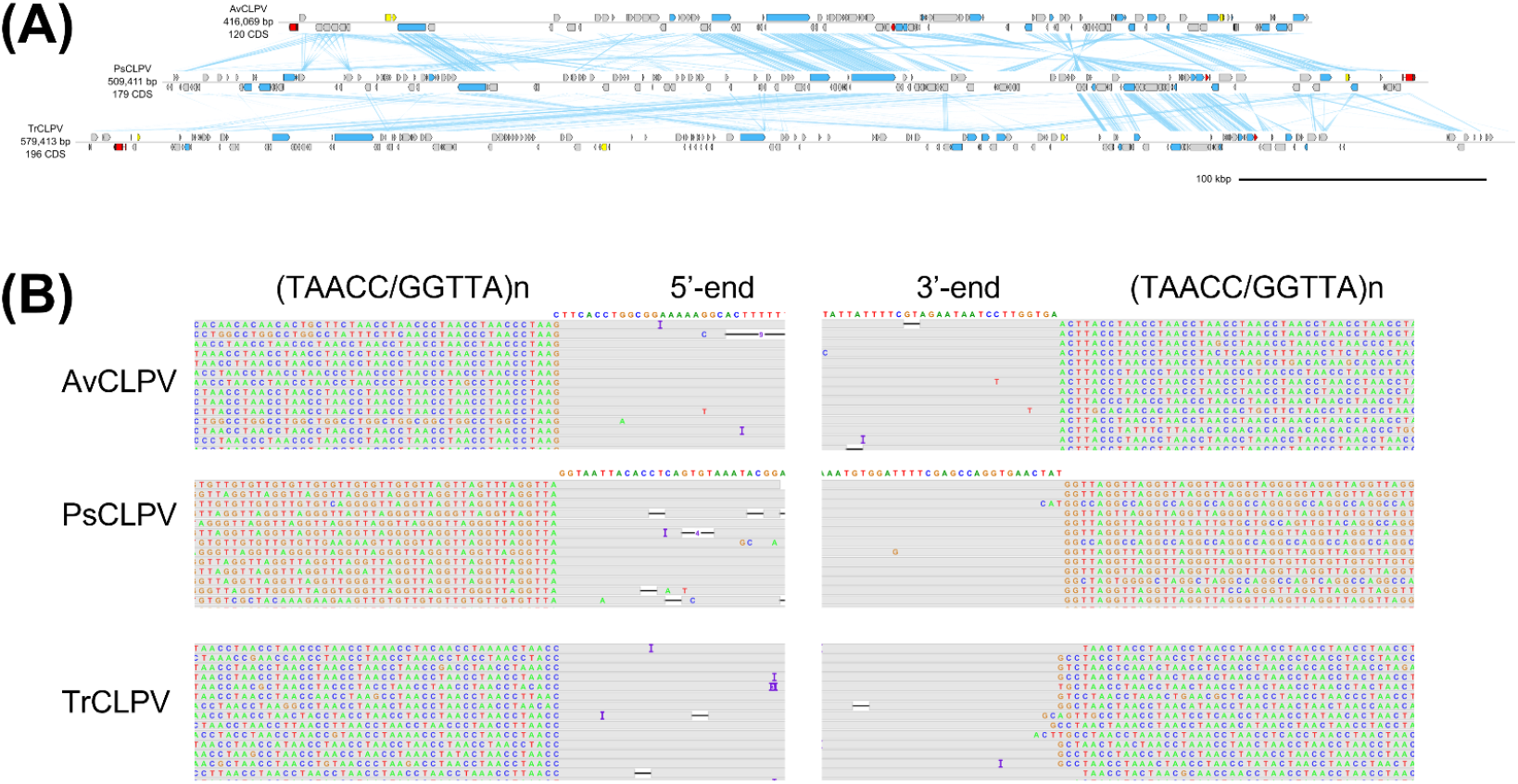
The clopoviruses. (A) Genome diagrams of clopoviruses. Arrows indicate predicted genes and their transcriptional orientations; blue, WSSV homologs; yellow, baculoviral inhibitor of apoptosis repeat-containing proteins; gray: hypothetical and other eukaryotic-like proteins; red, tyrosine recombinases. Blue ribbons indicate pairwise TBLASTX hits (e-value :1-e3, bitscore: 50) (B) ONT read alignments flanking the 5’- and 3’-ends of clopovirus genomes.

**Figure 6.**
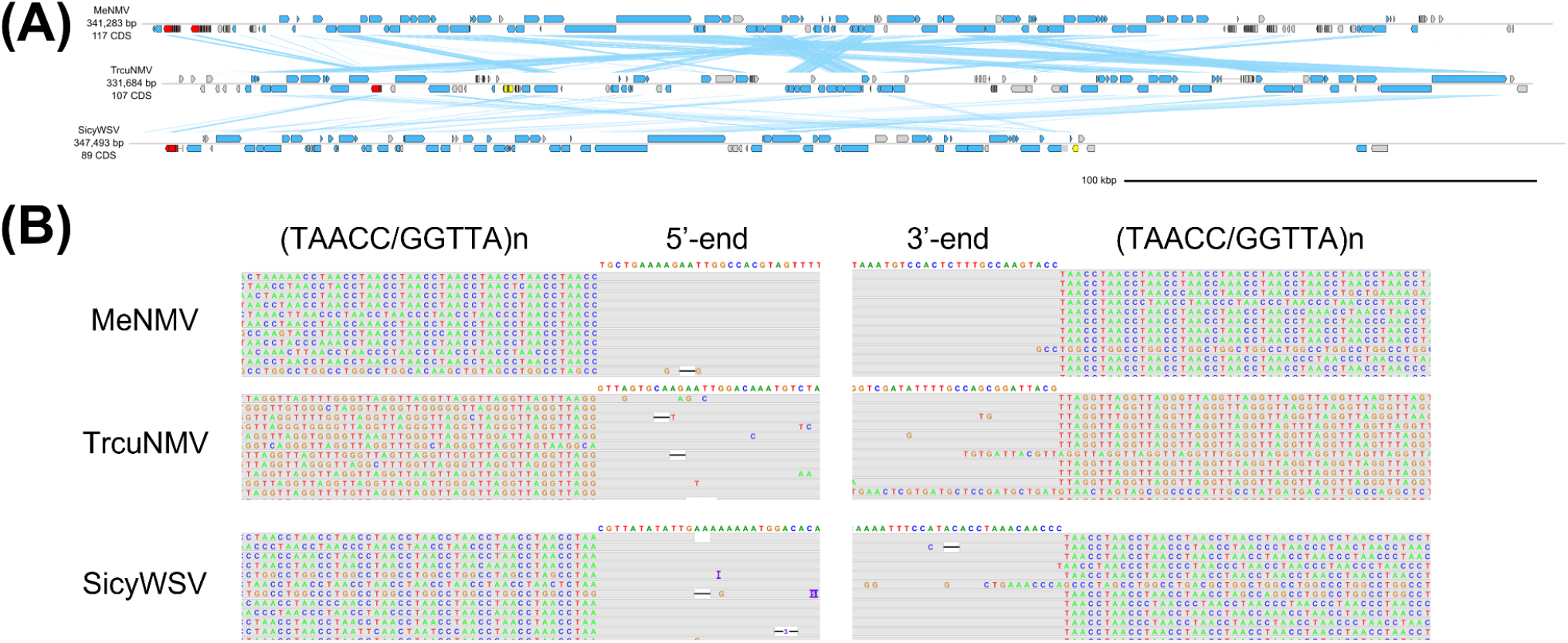
Telomere-inserting whispoviruses. (A) Genome diagrams of whispoviruses. Arrows indicate predicted genes and their transcriptional orientations; blue, WSSV homologs; yellow, baculoviral inhibitor of apoptosis repeat-containing proteins; gray: hypothetical and other eukaryotic-like proteins; red, tyrosine recombinases. Blue ribbons indicate pairwise TBLASTX hits (e-value :1-e3, bitscore: 50) (B) ONT read alignments flanking the 5’- and 3’-ends of whispovirus genomes.

All clopovirus genomes contained a stretch of (GGTTA/TAACC)n repeats, suggesting that clopoviruses specifically insert into this sequence motif (Figure 5B). However, most ONT reads mapping to these regions span the (GGTTA/TAACC)n repeat to align to either end, suggesting that many of the clopovirus copies exist as episomes or concatemers. For consistency with other nimaviral MAGs, we removed (GGTTA/TAACC)n from the clopoviral genome assemblies to produce linear contigs.

Together, these results reveal the presence of a divergent nimavirus lineage in terrestrial isopods, which we call clopoviruses. Clopoviruses possessed 19 ancestral nimaviral genes, of which 10 are core genes (Table 3). Given the small number of genes shared with other nimaviruses, clopoviruses could be classified into a novel family.

### Whispoviruses

We believe that the remaining nimaviruses can be united under *Whispovirus*, the only genus currently recognized by ICTV, due to their coherent phylogenetic clustering (Figure 1).

### Telomere-associated tyrosine recombinases

All telomere-associated nimaviruses shared a distinct family of tyrosine recombinase (YR), a site-specific DNA recombinase that is pervasive in prokaryotes but are rarely documented in eukaryotes (19, 20) (21) (Figure 7). The majaniviral YRs are arm-binding domain-containing tyrosine recombinases (22) that typically contained a C2H2 zinc binding domain and a N-terminal SAM-like domain (Figure 7A). We located five out of seven key residues important for YR functions (19, 23). The telomere-associated YRs are distinct from YRs encoded by nudiviruses (24) or NCLDV (25), nor are they closely related to known eukaryotic YR elements such as DIRS (26), Enterprise, and Cryptons (27, 28) (Figure 7B, C).

**Figure 7.**
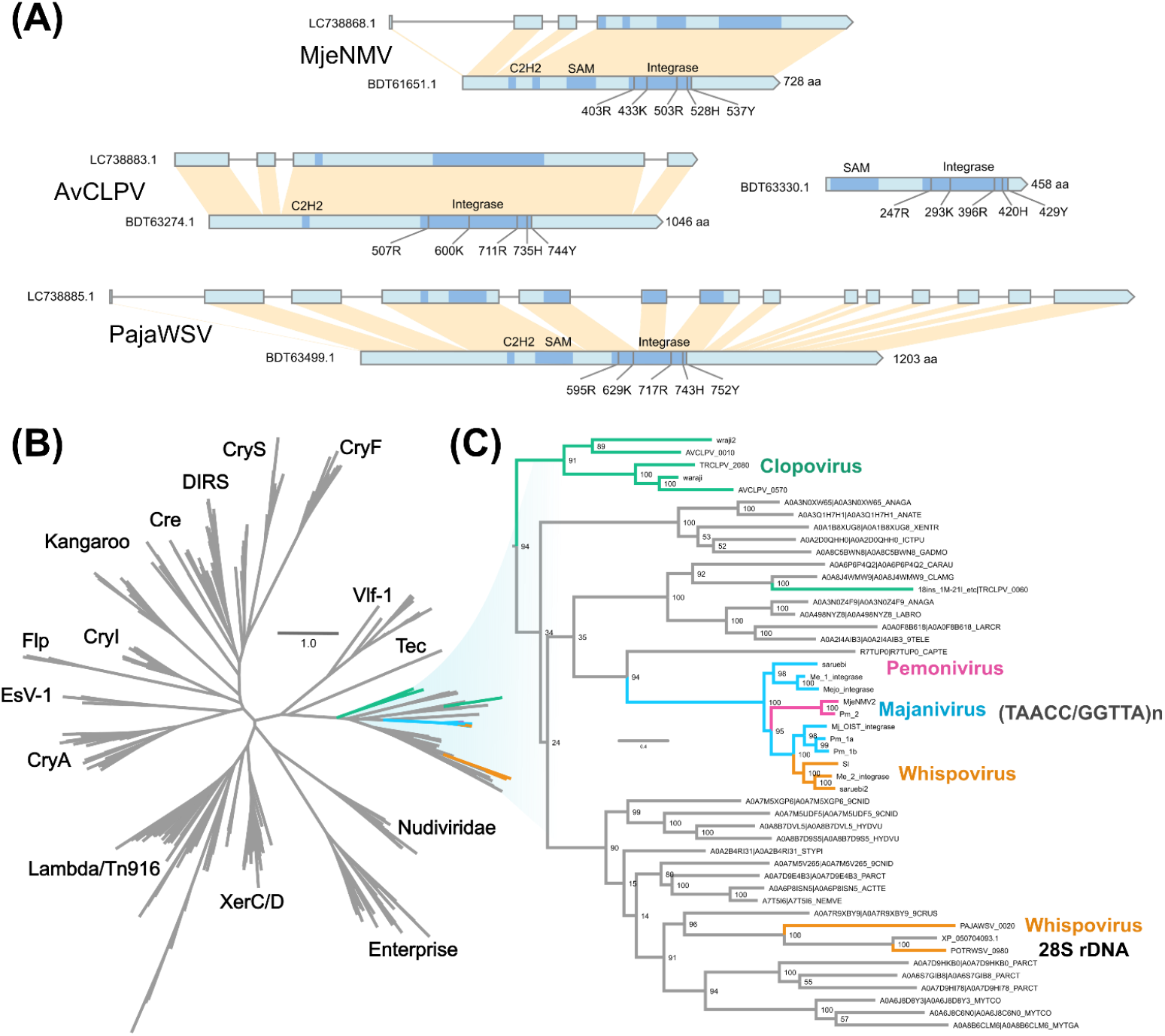
Nimaviral tyrosine recombinases. (A) Gene diagram of nimaviral tyrosine recombinases. (B) Maximum-likelihood phylogenetic tree of 342 tyrosine recombinases (555 sites; model: LG+R7). (C) Subtree of (B) showing the phylogenetic relationships of the nimaviral tyrosine recombinases.

### 28S rDNA-associated tyrosine recombinase in the closest WSSV relatives

Portunus trituberculatus whispovirus (PotrWSV; Supplementary Files 3 and 4) and Pasiphaea japonica whispovirus (PajaWSV) are the closest relatives of WSSV analyzed in this study, forming stem groups leading to WSSV (Figure 1, Figure 8). PortWSV was discovered from the genome sequencing data of the swimming crab *Portunus trituberculatus* (14). PajaWSV, identified from the shotgun sequence data of the Japanese glass shrimp (*Pasiphaea japonica*), is the closest relative of WSSV characterized in this study. PajaWSV and WSSV shares an average amino acid identity of 42.94%. PajaWSV and PortWSV insert into the host 28S rDNA with a 11-mer target site duplication (5’-CCGTCGCGRGAC-3’), a conserved motif occurring within 28S rDNA (Figure 8B and C).

**Figure 8.**
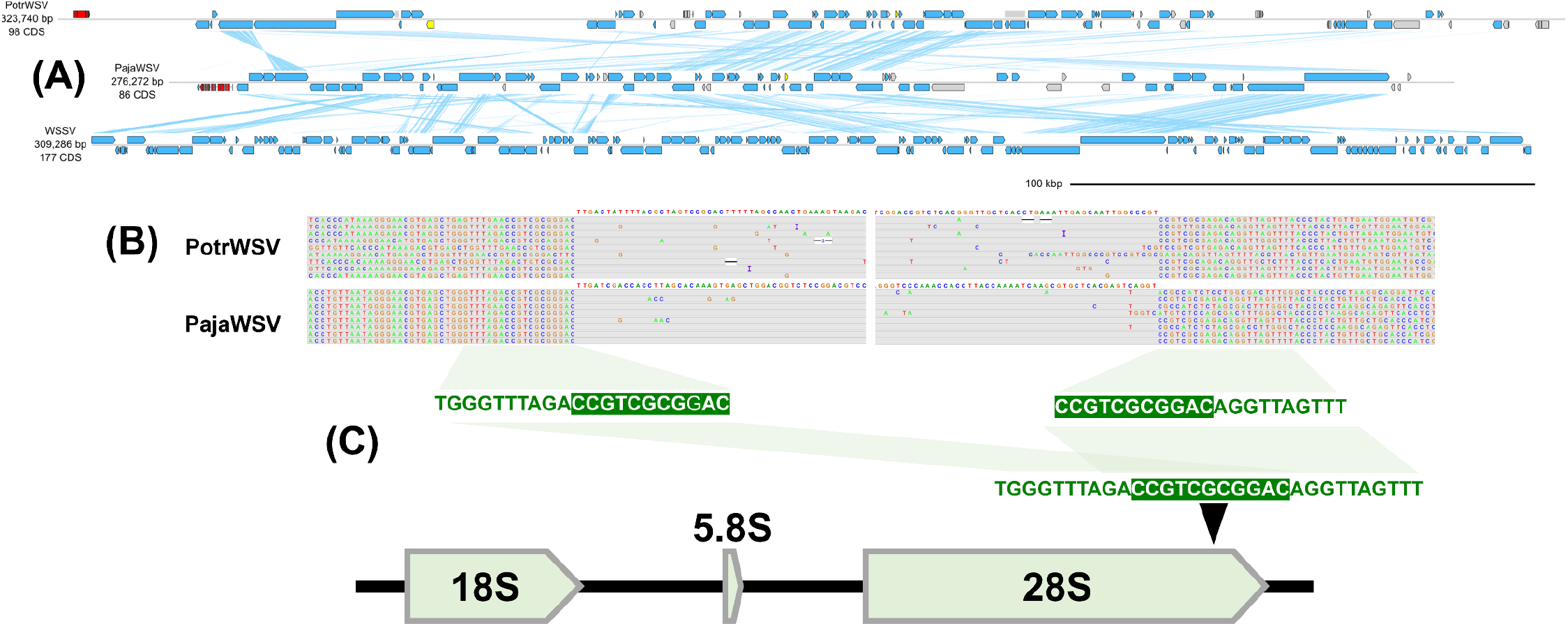
Genome diagrams of 28S rDNA-specific whispoviruses. (A) Genome diagram of sesarmid nimavirus genomes. Arrows indicate predicted genes and their transcriptional orientations; blue, WSSV homologs; yellow, baculoviral inhibitor of apoptosis repeat-containing proteins; gray: hypothetical and other eukaryotic-like proteins; red, tyrosine recombinases. Blue ribbons indicate pairwise TBLASTX hits (e-value :1-e3, bitscore: 50) (B) ONT read alignments flanking the 5’- and 3’-ends of 28S rDNA-specific whispoviruses. (C) 28S rDNA-specific whispovirus target site duplication.

PajaWSV and PotrWSV shared predicted multi-exon tyrosine recombinase genes that are phylogenetically related to telomere-associated YRs (Figure 7). BLASTP search revealed additional YR-like proteins from decapod crustaceans, although they were not associated with nimaviruses. Inclusion of the additional YRs into the phylogenetic tree led to the conclusion that PajaWSV and PotrWSV YRs were not immediate phylogenetic neighbors, raising the possibility that the YR genes in the two virus genomes were acquired independently. Collectively, these results suggest that two immediate WSSV relatives employ a distinct family of tyrosine recombinase to integrate into host 28S rDNA, although whether the tyrosine recombinase genes are orthologous remains an open question.

### Ginger2 exaptation in sesarmid nimaviruses

Sesarmid crabs *Orisarma intermedium* (formerly known as *Sesarmops intermedium*) and *Orisarma dehaani* (formerly known as *Chiromantes dehaani*) harbor endogenous nimavirus genomes (Figure 9) (10). The genome sequences of Seasrmops intermedium nimavirus (SiNMV; LC738884.1) and Chiromantes dehaani nimavirus (CdNMV) were assembled into sequences of 265-kb and 285-kb, respectively. The two sesarmid nimavirus genomes are extensively colinear and share 94% average nucleotide identity. Both viruses insert into (AT/TA)n repeats and are flanked by a 46-base teminal inverted repeat (TIR: 5’-GTTGTGCCTAATAAGGATAATGACTCATTAACGCTAATAGGTAACG-3’). The presence of a clearly defined inverted repeats was unique to the sesarmid nimaviruses.

**Figure 9.**
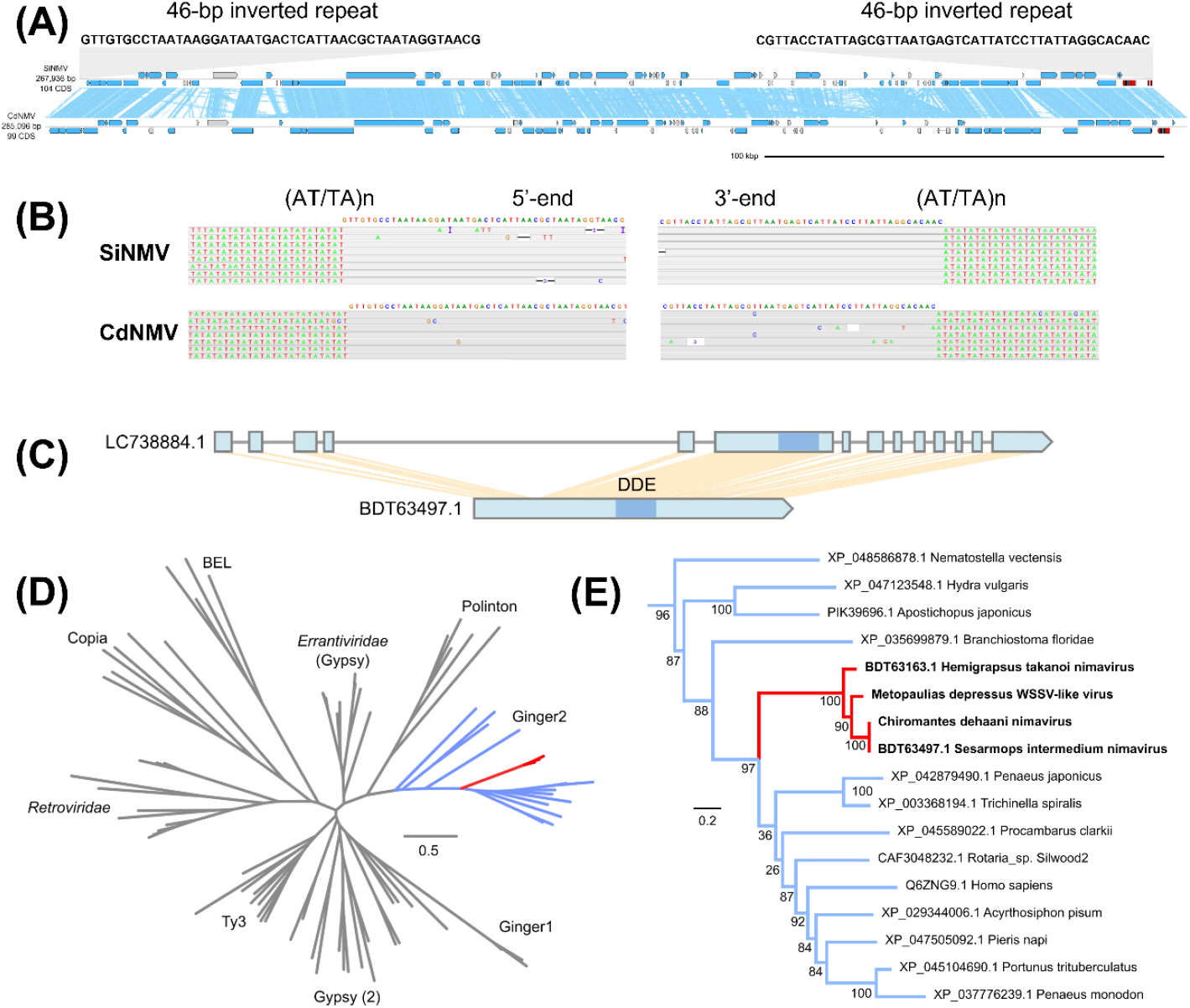
Ginger2 exaptation in sesarmid nimaviruses. (A) Genome diagram of sesarmid nimavirus genomes. Arrows indicate predicted genes and their transcriptional orientations; blue, WSSV homologs; gray: hypothetical and eukaryotic-like proteins; red, intgrases. Blue ribbons indicate pairwise TBLASTX hits (e-value :1-e3, bitscore: 50) (B) 5’- and 3’-end flanking sequences of the SiNMV genome. (C) Gene diagram of the integrase-like gene in the SiNMV genome. (D) Maximum-likelihood phylogenetic tree of 57 DDE transposases (422 sites; model: Q.insect+I+G4). (E) Subtree of (D) showing the phylogenetic relationships of Ginger2-like elements.

SiNMV and CdNMV genomes lacked YRs but possessed intron-containing genes with structural similarities to retroviral integrases (Figure 9) (29). These nimaviral integrases clustered with Ginger2 transposable elements, a group of intron-containing cut-and-paste transposable element with TIRs (30). Hemigrapsus takanoi nimavirus also lacked YR and had a similar integrase gene (Figure 9), Along with the presence of 46-bp TIR flanking the sesarmid nimavirus genomes, these results suggest that sesarmid endogenous nimaviruses, and possibly Hemigrapsus takanoi nimavirus, have adapted a distinct family of integrase-like genes for AT/TA-motif-specific integrations.

## Discussion

It has long been known that crustacean genomes harbor various WSSV-like sequences (7–9, 31), but the reasons why they are present has remained unknown. The present results demonstrate that endogenous nimaviruses selectively insert into specific genomic contexts, and this specificity is correlated with the types of integrases they encode. We propose that endogenous nimaviruses are selfish genetic elements that persist within the host genomes (32), and that the capture of integrase genes with different insertion specificities has allowed nimaviruses to persist as genomic parasites colonizing different repetitive motifs representing genomic niches (33, 34). We note that these endogenous nimaviruses are distinct from fragmented viral insertions that may produce potentially immunogenic transcripts (35–37). The selfish nature of transposable elements could explain the persistence of endogenous nimaviruses even without a perceivable selective advantage to the host (38).

While it is possible that promiscuous integration followed by biased selective retention produced the appearance of selective integration, it is unlikely that only one of a widevariety of repetitive motifs present in the host genome would be tolerated. Therefore, the most likely explanation for the observed insertion selectivity is that it is mediated by site-specific integrases.

Overall, the distribution of integrases among nimaviruses does not strictly align with their phylogenetic relationships, indicating that nimaviruses have acquired integrases multiple times through evolutionary history. Interestingly, we observed that nimaviruses from phylogenetically distinct lineages, such as sesarmid nimaviruses and Hemigrapsus takanoi nimavirus, can possess mutually similar integrases, which raises the possibility that integrase genes may have been shared among different lineages of nimaviruses (39).

Integration of naldaviruses into host genomes has occurred many times among naldaviruses (24, 40–44),, and some, such as polydnaviruses, have even become domesticated to serve host functions (24, 45–47). Endogenization is also prevalent among herpesviruses, with the *Teratorn* element in medaka fish being a well-known example (48–51). Overall, it appears that nuclear double-stranded DNA viruses have evolved along a spectrum between endogenous and exogenous states, swaying between the two extremes.

Repeat-specific integration is believed to be a survival strategy employed by transposable elements in order to minimize negative effects on host fitness (52–54). The prevalence of telomere-repeat-specific nimaviruses in penaeid shrimps and terrestrial isopods may be due to a combination of this viral survival strategy and the abundance of simple sequence repeats, including telomere-like repeats (55), in the genomes of these organisms (15, 56–59).

Endogenous nimaviruses tenaciously retain structural protein genes, suggesting that they maintain the capability to form viral particles and transmit between hosts. We speculate that endogenous nimaviruses in crustacean genomes are analogous to prophages in bacterial genomes, which can remain dormant until certain stressors trigger their reactivation. Conservation of PIFs among endogenous nimaviruses is particularly noteworthy (16), as they may facilitate the oral transmission of viral particles through cannibalism of dead hosts, which is a common transmission route of WSSV (60–62).

Our analyses may be biased by incomplete taxon sampling of the host and the scarcity of exogenous nimavirus genomes. However, the lack of observed diversity could also reflect the actual rarity of exogenous nimaviruses circulating in the environment. To date, we have not been able to identify nimavirus-like sequences in environmental metagenomes, and despite the long history of modern shrimp aquaculture, WSSV remains the only pathogenic nimavirus of penaeid shrimps. We hypothesize that exogenous nimaviruses are rare and the emergence of a pathogenic nimavirus is an even more unusual event. To confirm this hypothesis, it would be valuable to conduct thorough metagenomic surveys, similar to one conducted in *Drosophila melanogaster* (63), to assess the prevalence of exogenous nimaviruses and other double-stranded DNA viruses in various crustacean species.

## Materials and Methods

### General sequencing, assembly, and annotation strategy

Decapod crustacean genomes are gigabase-sized and extremely rich in repetitive elements, making whole genome assembly challenging (7, 8, 56, 57, 64–66). To circumvent this difficulty, we performed shallow-depth genome survey sequencing and assembled the reads as a metagenome and picked up viral contigs. We also analyzed publicly available genomic sequences where available (14, 15).

### Crustacean genome survey sequencing

We sequenced a total of 17 crustacean genomes using Illumina and Oxford Nanopore Technologies (ONT) platforms. Genomic DNA was extracted from muscle or whole animal using phenol-chloroform-isoamyl alcohol extraction or MagAttract HMW DNA Kit (Qiagen), and further purified using NucleoBond columns (Machery Nagel). For some samples, genomic DNA was size-selected using a custom PEG/NaCl precipitation buffer [9% PEG8000 (w/v), 1M NaCl, 10 mM Tris-HCl (pH 8.0)] (67). For four of the samples (*Litopenaeus vannamei, Hemigrapsus takanoi, Sesarmops intermedium*, and *Chiromantes dehaani*) we used gDNA preparations generated in a previous study (10).

ONT long-read libraries were prepared using the Ligation Sequencing Kit (SQK-LSK109, ONT), NEBNext Companion Module for Oxford Nanopore Technologies Ligation Sequencing (E7180, NEB), and Agencourt AMPure XP beads (Beckman Coulter). Libraries were size selected using the PEG/NaCl precipitation buffer described above. The ONT libraries were sequenced on R9.4.1 flow cells, with multiple nuclease flush (EXP-WSH004, ONT) and priming (EXP-FLP002, ONT) before library loading. The fast5 files were base-called using Guppy v5.0.11, v5.0.13, or v5.0.16, with the super accuracy mode. The fast5 files of the previously published *M. japonius* ONT reads (Ginoza2017; BioSample Accession No. SAMD00276454) (57) were base-called using Guppy v5 v5.0.13 with super accuracy mode.

Library preparation and sequencing on Illumina HiSeq 4000 (2×150-bp) was carried out by Eurofins Genomics (Tokyo). Raw Illumina reads were quality trimmed by Fastp v0.20.1(68). The filtered Illumina reads were also used for *de novo* assembly and polishing.

### Publicly available datasets

Publicly available whole genome shotgun sequence data of *Portunus trituberculatus* (14) and *Trachelipus rhatkii* (15) were downloaded from the NCBI database. The raw reads were analyzed in a similar manner to other crustacean genome data.

### *De novo* assembly of shotgun sequence data and virus discovery

The filtered Illumina reads were *de novo* assembled using SPAdes (69). Combinations of SPAdes versions and parameters varied depending on the time of the analysis, sequencing coverage, and data complexity. The SPAdes assemblies were used to salvage low-copy nimaviral sequences that could not be fully recovered from ONT assemblies, including Nima-1_Lva and SiNMV.

The ONT reads were filtered by 5-, 10-, or 20-kb length cutoffs using SeqKit (70) and were *de novo* assembled by Flye v. 2.9 (71) in metagenome mode. The primary ONT assemblies were visualized by Bandage v0.8.1 (72) and screened for nimaviral sequences were by TBLASTN searches querying WSSV proteins. The identified nimaviral contigs were used as the bait to map back the ONT reads by Minimap2 (73), and the mapped ONT reads were reassembled by Flye v2.9 (71) in normal mode. This generated consensus nimaviral genome sequences that we believe are close representations of the original viral genomes.

The contigs were subjected to multiple rounds of polishing involving Medaka v1.4.3, HyPo v1.0.3 (74), Pilon v1.24 (75), and/or POLCA (76), using ONT reads and Illumina reads. The actual combinations of polishers differ between the viral genomes. ONT and Illumina reads were mapped backed by Minimap2 and visualized using IGV to inspect read coverage and misassemblies (77). Assembly errors were manually curated.

### Gene prediction and annotation

Endogenous nimaviruses contained eukaryotic-like genes with introns, which cannot be predicted by prokaryotic gene prediction programs. To recover both classes of protein-coding genes, we used different gene prediction programs and integrated the outputs into a nonredundant annotation. Open reading frames were predicted by Prodigal v2.6.3 (78), and the predicted proteins were queried against the proteomes of nimaviruses (WSSV, Marsupenaeus japonicus endogenous nimavirus, Penaeus monodon endogenous nimavirus, Hemigrapsus takanoi nimavirus, Metapenaeus ensis nimavirus, Sesarmops intermedium nimavirus; last accessed December 2021) and arthropods (*Marsupenaeus japonicus, Litopenaeus vannamei, Penaeus monodon, Portunus trituberculatus*, and *Homarus americanus*; last accessed December 2021) (79), using BLASTP. The BLASTP output were merged by Automated Assignment of Human Readable Descriptions (AHRD) pipeline (80) into a table containing functional description. The genomic coordinates corresponding to nimaviral-like proteins were masked by BEDtools (81), and the remaining coordinates were forwarded to *ab initio* eukaryotic-like gene prediction by Augustus v3.3.3 (82) using the *Apis mellifera* gene model (11). The use of *Apis mellifera* gene model was inspired by Bao et al. (2020) (11). The predicted proteins (generated by gffread (83)) were BLASTP searched against the abovementioned nimaviral and arthropod proteomes, and the BLASTP output were passed to AHRD to generate final functional annotations. The GFF3 annotation files were converted into DDBJ flat files using GFF3toDDBJ (https://github.com/yamaton/gff3toddbj) and FFconv (https://www.ddbj.nig.ac.jp/ddbj/ume-e.html).

### Comparative genomic analysis and visualization

Genome diagrams were generated by a custom script (https://github.com/satoshikawato/bio_small_scripts/blob/main/plot_linear_genome.py). Average nucleotide identity (ANI) and average amino acid identity (AAI) values were calculated by the ANI calculator (http://enve-omics.ce.gatech.edu/ani/) and the AAI calculator (http://enve-omics.ce.gatech.edu/aai/), respectively (84–86).

### Phylogenomic analysis

Amino acid sequences of nine nimaviral core genes (wsv026, wsv282, wsv289, wsv303, wsv343, wsv360, wsv433, wsv447, and wsv514; 13,007 amino acids) were aligned using MAFFT v7.505 (87), and the alignments were trimmed using trimAl v1.2 (88). Maximum likelihood phylogenetic analysis was performed using IQ-TREE2 v2.2.0.3 (89).

### Analysis of integrase genes

Multiple sequence alignments of representative tyrosine recombinase families (*CryA, CryF, CryI, CryS, Kangaroo*, and *Tec*) were downloaded from Kojima et al.(28) and queried against UniRef30 2020 Februrary version on the HHblits server (https://toolkit.tuebingen.mpg.de/tools/hhblits; last accessed September 13, 2022) (90). Cre recombinase, Enterprise, VLF1, XerCD, Tec, Tn916, and Lambda integrase homologs were prepared by querying individual proteins against UniRef30 Feb. 2022 by HHblits or NCBI non-redundant protein database by BLASTP. The proteins were aligned on MAFFT server (last accessed September 13, 2022) (87) with default settings. The alignments were then iteratively refined using CD-HIT (75% identity cutoff) (91) and MaxAlign (92) implemented on the MAFFT server. The alignments were subjected to HHpred server to identify regions exhibiting similarities to YR domains, cropped, aligned by MAFFT and further refined. The resulting YR entries were merged into a single alignment by MAFFT.

## Acknowledgements

This research was supported by Grants-in-Aid for Scientific Research from the Japan Society for Promotion of Science (JSPS) (JSPS KAKENHI Grant Numbers JP15H02462, JP19H00949 and 19J21518) and by Science and Technology Research Partnership for Sustainable Development from the Japan Science and Technology Agency (SATREPS Grant Number JPMJSA1806).

## Data availability

The raw reads generated in this study are deposited to DDBJ/NCBI/ENA database under the BioProject ID PRJDB13888. The accession numbers of the nimaviral MAG assemblies are provided in Table 2. TrCLPV genome and protein sequences are available as Supplementary Files 1 and 2, respectively. PotrWSV genome and protein sequences are available as Supplementary Files 3 and 4, respectively. Supplementary Files 1-4 are available on FigShare (https://figshare.com/s/bf24bce327dff99ee0b0).

