## Supplemental Figure 1 for "Integrase-associated niche differentiation of endogenous large DNA viruses in crustaceans"

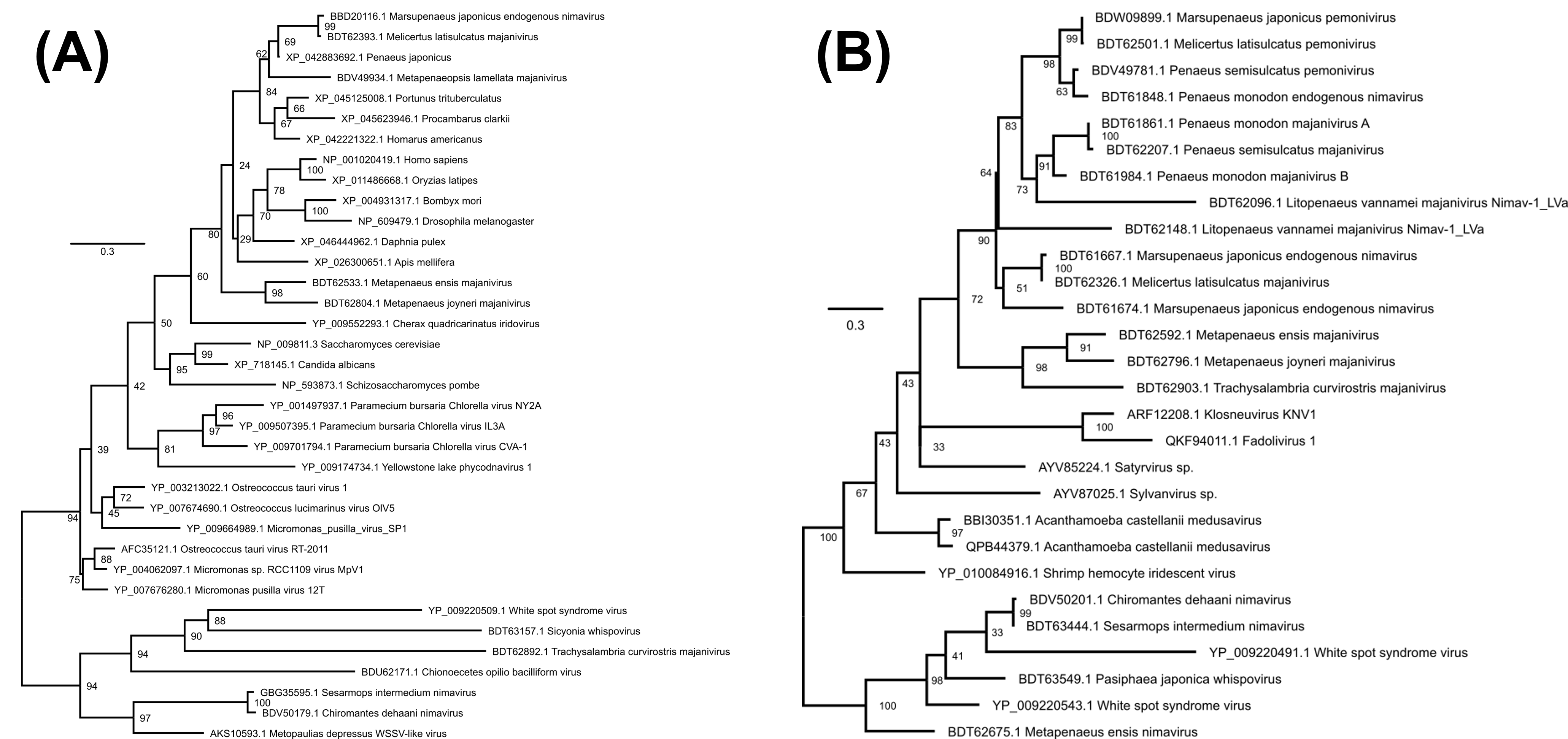

### Supplementary Figure 1. Maximum phylogenetic trees of wsv112 and wsv206-like proteins

(A) Maximum-likelihood phylogenetic tree of wsv112 (dUTPase)-like proteins (112 sites; model: LG+G4).
(B) Maximum-likelihood phylogenetic tree of wsv206-like proteins (111 sites; model: LG+G4).
